# Spatio-temporal distribution of *Spiroplasma* infections in the tsetse fly (*Glossina fuscipes fuscipes*) in northern Uganda

**DOI:** 10.1101/591321

**Authors:** Daniela I. Schneider, Norah Saarman, Maria G. Onyango, Chaz Hyseni, Robert Opiro, Richard Echodu, Michelle O’Neill, Danielle Bloch, Aurélien Vigneron, T.J. Johnson, Kirstin Dion, Brian L. Weiss, Elizabeth Opiyo, Adalgisa Caccone, Serap Aksoy

## Abstract

Tsetse flies (*Glossina* spp.) are vectors of parasitic trypanosomes, which cause human (HAT) and animal African trypanosomiasis (AAT) in sub-Saharan Africa. In Uganda, *Glossina fuscipes fuscipes* (*Gff*) is the main vector of HAT, where it transmits Gambiense disease in the northwest and Rhodesiense disease in central, southeast and western regions. Endosymbionts can influence transmission efficiency of parasites through their insect vectors via conferring a protective effect against the parasite. It is known that the bacterium *Spiroplasma* is capable of protecting its *Drosophila* host from infection with a parasitic nematode. This endosymbiont can also impact its host’s population structure via altering host reproductive traits. Here, we used field collections across 26 different *Gff* sampling sites in northern and western Uganda to investigate the association of *Spiroplasma* with geographic origin, seasonal conditions, *Gff* genetic background and sex, and trypanosome infection status. We also investigated the influence of *Spiroplasma* on *Gff* vector competence to trypanosome infections under laboratory conditions.

Generalized linear models (GLM) showed that *Spiroplasma* probability was correlated with the geographic origin of *Gff* host and with the season of collection, with higher prevalence found in flies within the Albert Nile (0.42 vs 0.16) and Achwa River (0.36 vs 0.08) watersheds and with higher prevalence detected in flies collected in the intermediate than wet season. In contrast, there was no significant correlation of *Spiroplasma* prevalence with *Gff* host genetic background or sex once geographic origin was accounted for in generalized linear models. Additionally, we found a potential negative correlation of *Spiroplasma* with trypanosome infection, with only 2% of *Spiroplasma* infected flies harboring trypanosome co-infections. We also found that in a laboratory line of *Gff*, parasitic trypanosomes are less likely to colonize the midgut in individuals that harbor *Spiroplasma* infection. These results indicate that *Spiroplasma* infections in tsetse may be maintained by not only maternal but also via horizontal transmission routes, and *Spiroplasma* infections may also have important effects on trypanosome transmission efficiency of the host tsetse. Potential functional effects of *Spiroplasma* infection in *Gff* could have impacts on vector control approaches to reduce trypanosome infections.

**Author Summary:** We investigated the association of symbiotic *Spiroplasma* with the tsetse fly host *Glossina fuscipes fuscipes* (*Gff*) to assess if *Spiroplasma* infections are correlated with *Gff* genetic background, geography, or season and its interaction with trypanosome parasites. We analyzed distribution and prevalence of *Spiroplasma* infections across different *Gff* sampling sites in northern and western Uganda, and found that the symbiont is unevenly distributed and infections have not reached fixation within these sampling sites. We tested for associations with geographic origin of the collections, seasonal environmental conditions at the time of collection, *Gff* host genetic background and sex, plus trypanosome co-infections. *Spiroplasma* prevalence was strongly correlated with geographic origin and seasonal environmental conditions. Our parasite infection data suggested a negative correlation of *Spiroplasma* with trypanosome infection, with only 5 out of 243 flies harboring trypanosome co-infections. We further investigated the influence of *Spiroplasma* on trypanosome parasite infections in the laboratory. We found that trypanosomes were less likely to establish an infection in *Gff* individuals that carried *Spiroplasma* infections. Our results provide new information on host-endosymbiont dynamics in an important human disease vector, and provide evidence that *Spiroplasma* may confer partial resistance to *Gff* trypanosome infections. These findings provide preliminary evidence that a symbiont-based control method could be successful in combating tsetse trypanosome transmission to humans and livestock in sub-Saharan Africa.

## Introduction

Tsetse flies (*Glossina spp*.) are vectors of parasitic African trypanosomes that cause human African trypanosomiasis (HAT, commonly referred to as sleeping sickness) and African animal trypanosomiasis (AAT, also known as Nagana in cattle) [1-3]. Several major HAT epidemics in sub-Saharan Africa have occurred during the last century, with the most recent one resulting in over half a million deaths in the 1980s [4-7]. An ambitious campaign led by WHO and international partners has now reduced the prevalence of HAT in west Africa [8] to a threshold considered irrelevant for epidemiological considerations, but millions continue to live at risk of contracting HAT in tsetse inhabited areas [9-11]. Despite calls for elimination of HAT by 2030, there is a lack of effective tools for long-term control of the disease (e.g., vaccines and field-ready diagnostic assays). Furthermore, the presence of animal reservoirs threatens disease elimination efforts going forward, particularly in East Africa (reviewed in [12]) and necessitates the inclusion of vector control applications. Practical interventions [13-15], as well as mathematical models [16-18], suggest that vector control can accelerate efforts for reaching the disease elimination phase. Thus, enhancing the vector-control tool box with effective and affordable methods is a desirable goal. Biological approaches that reduce vector reproduction as well as vector competency have emerged as promising means to reduce disease transmission [19].

Variation in the microbiota associated with tsetse flies can influence their pathogen transmission dynamics [20-24]. Several symbiotic microorganisms have been described from laboratory and field populations of tsetse, including the obligate *Wigglesworthia glossinidia*, commensal *Sodalis glossinidius*, parasitic *Wolbachia* and more recently *Spiroplasma* [25-31]. A survey of Ugandan *Gff* revealed that all individuals harbored *Wigglesworthia,* while *Sodalis* and *Wolbachia* associations were sporadic [28,32,33]. *Spiroplasma* was found in the tsetse species that belong to the *Palpalis* group (*Gff, G. p. palpalis* and *G. tachinoides*), while the tsetse species in the other two subgroups, *Morsitans* and *Fusca*, lacked associations with this microbe [31].

The bacterium *Spiroplasma* confers protection against nematodes, fungi and parasitoid wasps in several insects [34-36]. *Spiroplasma* also acts as a reproductive parasite that induces a male-killing effect in some arthropod hosts [37-40]. Based on phylogenetic analyses, the *Spiroplasma* species infecting *Gff* (both field-caught individuals and one laboratory line) is most closely related to the *Citri*-*Chrysopicola*-*Mirum* clade that confers protection against parasitoid wasps, nematodes and fungal pathogens in the fruit fly and aphid hosts [31]. In addition, among the *Spiroplasma* species within this clade are some that are pathogenic to plants and invertebrates and some that exhibit a male killing phenotype in ladybirds, fruit flies, and some butterfly species they infect (reviewed in [41]). *Spiroplasma* function(s) within the tsetse host remain unknown.

In Uganda, *Gff* is the main vector of HAT, where it transmits the chronic Gambiense disease caused by *Trypanosoma brucei gambiense* in the northwest and acute Rhodesiense disease caused by *T. b. rhodesiense* in the center, southeast and west (reviewed in [42]). It is possible that differences in *Gff* trypanosome susceptibility (vector competence) among varying geographic regions could be influenced by *Spiroplasma*, but patterns of *Spiroplasma* occurrence remain unexplored. *Spiroplasma* infection success may be influenced by seasonal fluctuations in host *Gff* population health (fitness) and density. Seasonal changes in the environment can dramatically alter host-endosymbiont dynamics through changes in *Gff* host physiological status (e.g. hemolymph lipid levels [43,44]) and tsetse population density [45,46]. Collections of *Gff* in Uganda during different seasons provides the opportunity to test for influence of seasonal fluctuations on *Spiroplasma* prevalence.

Additionally, patterns of *Spiroplasma* prevalence are likely influenced by *Gff* host genetic background, either because of vertical transmission that follows host inheritance patterns, or because of lineage-specific coevolutionary dynamics. *Gff* in Uganda has high genetic structuring at multiple spatial scales, which provides the opportunity to test for influence of *Gff* genetic background on *Spiroplasma*. There are three distinct *Gff* genetic clusters in the north, west and south of the country, and further population structure that separates the northwest from the northeast with ongoing gene flow between them in an admixture zone [47-49]. For this study, we focus our work on the north and west of Uganda in five watersheds. This sampling included four nuclear (biparentally inherited) genetic backgrounds: the northwest genetic unit (NWGU), the northeast genetic unit (NEGU), the west genetic unit (WGU) and the admixed (ADMX) genetic background intermediate to the NWGU and NEGU [47-49]. This sampling also included three mitochondrial (maternally inherited) genetic backgrounds: a group of related haplotypes associated with the NWGU known at “haplogroup A” (mtA), a group of related haplotypes associated with the NEGU known at “haplogroup B” (mtB), and a group of related haplotypes associated with the WGU known at “haplogroup C” (mtC) [47-49].

In this study, we *(i)* assessed the infection prevalence of *Spiroplasma* in *Gff* among five watersheds in northern and western Uganda, *(ii*) tested the effect of seasonal environmental variations, host *Gff* genetic background and sex, and trypanosome co-infections on *Spiroplasma* prevalence, and *(iii)* investigated the influence of *Spiroplasma* infections on *Gff* trypanosome transmission ability under laboratory conditions. We discuss how our results can elucidate potential functional associations of *Spiroplasma* with its tsetse host, and the potential applications of this knowledge to disease control.

## Methods

### Field sampling

A total of 1415 *Gff* individuals collected from northern and western Uganda during the period of 2014-2018 across the wet (April - May and August - October), intermediate (June - July), and dry (December - March) seasons were assayed for *Spiroplasma* infection (S1 Table). Flies were collected on public land using biconical traps. Samples included in this study were chosen to represent a wide range of environmental conditions and *Gff* backgrounds (Fig 1).). The location of each sampling site was placed on a map of Uganda using QGIS v2.12.1 (August 2017; http://qgis.osgeo.org) with free and publicly available data from DIVA-GIS (August 2017; http://www.diva-gis.org). We sampled 11 sites from the Albert Nile watershed (DUK, AIN, GAN, OSG, LEA, OLO, PAG, NGO, JIA, OKS, and GOR), where the majority of *Gff* belong to the NWGU/mtA genetic background, and a minority belong to the ADMX/mtB genetic background [47-49]. We sampled six sites from the Achwa River watershed (ORB, BOL, KTC, TUM, CHU, and KIL), where *Gff* belong to a mix of genetic backgrounds including NWGU/mtA, ADMX/mtA, ADMX/mtB, and NEGU/mtB [47-49]. We sampled six sites from the Okole River watershed (ACA, AKA, OCA, OD, APU, and UWA), where *Gff* hosts belong to a mix of genetic backgrounds including NWGU/mtA, ADMX/mtA, ADMX/mtB, ADMX/mtC, and NEGU/mtB [47-49]. We sampled two sites from the Lake Kyoga watershed (OCU and AMI), where the majority of *Gff* belong to the NEGU/mtB genetic background, and a minority belong to the ADMX/mtA genetic background [47-49]. Finally, we sampled a single site from the Kafu River watershed (KAF), where the majority of *Gff* belong to the WGU/mtC genetic background, and a minority belong to the WGU/mtA genetic background [47-49].

### Cloning and sequencing of *Spiroplasma 16S r*DNA and MLST genes

To assess the *Spiroplasma* strain infecting the field-collected samples, we cloned and sequenced Spiroplasma-*16S r*DNA and *rpo*B (RNA polymerase, subunit beta) from flies from two NGU sampling sites (GAN and GOR). The *16S r*DNA touchdown PCR was performed in 20 µL reactions containing 1× GoTaq® Green Mastermix (Promega, USA), 0.4 µM of each primer *Spir*RNAF and *Spir*RNAR (S2 Table) and 1 µL of template DNA (20-100ng). The PCR profile consisted of an initial denaturation for 3 mins. at 94°C, followed by 8 cycles of 1 min. at 94°C, 1 min. at 63°C, and 1 min. at 72 with a reduction in annealing temperature of 1°C/cycle (63-56°C). This step was followed by 30 cycles of 1 min. at 94°C, 1 min. at 55°C, and 1 min. at 72°C, with a final extension at 72°C for 10 min. Reaction set up for the *rpo*B-PCR was the same as for *16S r*DNA (S2 Table), and the PCR profile consisted of a 3-min. initial denaturation at 94°C, followed by 34 cycles of 90 sec. at 94°C, 90 sec. at 55°C, and 90 sec. at 72°C, with a final extension at 72°C for 10 min. All amplicons were gel purified using the Monarch® DNA Gel Extraction Kit (New England Biolabs, USA) and cloned into pGEM-T vector (Promega, USA). In total six *16S r*DNA and two *rpo*B clones were sequenced and analysed using the BLASTN algorithm and finally aligned using *Spiroplasma glossinidia* data from NCBI as reference.

To test for potential *Spiroplasma* strain variation among the different *Gff* sampling sites, we cloned and sequenced the Multi Locus Sequence Analysis (MLST) genes *dna*A, *fru*R, *par*E and *rpo*B from two individuals from ACA (Okole River) and OKS (Lake Kyoga) sampling sites. MLST-PCR reactions were performed with the same reaction set up as described above (S2 Table). The *par*E touchdown PCR was run as described above for *16S r*DNA but with 61°C/52°C annealing temperature. A touchdown PCR approach was also used for the *fru*R locus with 30 sec steps in each cycle and 60°C/52°C annealing temperature. Cloning and sequencing was performed as described above. All sequences were manually edited using Bioedit v7.1.9 [50] and aligned with related sequences available at NCBI using MUSCLE [51].

### *Spiroplasma, Wolbachia* and trypanosome infection status

At the time of collection, sex and wing fray information (all flies were wing fray 2-3, i.e. 4-8 weeks old) was recorded for each fly and midguts were dissected and microscopically analyzed for trypanosome infection status. The dissected guts and reproductive parts were stored in 95% ethanol for further analyses. Genomic DNA was extracted from female and male reproductive parts (RP, n = 1157) as well as from whole bodies (WB, n = 258) using DNeasy® Blood and Tissue Kit (Qiagen, Germany). We used flies from Pagirinya (PAG) for which we had DNA available from both the WB and RP tissues to compare the *Spiroplasma* infection prevalence to rule out potential DNA source bias in the prevalence results. We noted *Spiroplasma* infection of 18% in WB versus 17% in RP, which is not significantly different (Fisher’s *P* > 0.999). Consequently, both tissue datasets were pooled for the final infection prevalence analyses. The final sample set was composed of 893 females and 522 males.

*Spiroplasma* infection prevalence was determined by *16S r*DNA touchdown PCR as described above. The presence of *Wolbachia* was assessed by *wsp*-, *ARM*-and *VNTR*141-PCR amplifications. The *Wolbachia* Surface Protein gene (*wsp*) is a single copy membrane protein encoding gene, whereas *ARM* (*Wolbachia* A supergroup repeat motif) and *VNTR* (Variable number of tandem repeats) are high copy number repeat motives found in *Wolbachia*. The PCR reactions were performed as previously described in [52-54]. All amplified fragments were analysed on 1% agarose gels using Gel Doc EQ quantification analysis software (Bio-Rad, Image Lab™ Software Version 4.1).

### Assessment of variation in *Spiroplasma* density by qPCR

Potential *Spiroplasma* density differences between individuals and across seasons were tested via quantitative PCR (qPCR) using a CFX96 Real-Time PCR Detection System and iTaq™ Universal SYBR® Green Supermix (Biorad). The relative amount of *Spiroplasma* was calculated with the 2^-(ΔΔCt)^ method using primers targeting the *Spiroplasma* RNA polymerase beta gene (*rpo*B). Values were normalized against the single copy *Glossina* Peptidoglycan Recognition Protein-LA gene (*pgrp*-*la*). All primer sets are listed in S2 Table. The trypanosome infection status of each sample was assessed by microscopic analysis of dissected guts at the time of sample collection in the field.

### Mitochondrial DNA (mtDNA) sequencing and microsatellite genotyping

In addition to the samples typed for a previous study [47], we sequenced an additional 161 flies for a 491 bp fragment of mtDNA, and genotyped an additional 131 flies at 16 microsatellite loci. To do this, we extracted total genomic DNA from two legs of individual flies using the Qiagen DNeasy Blood & Tissue kit. For mtDNA sequencing, a 491 bp fragment from the cytochrome oxidase I and II gene (*COI, COII*) was amplified as described in [47]. PCR amplicons were run on 1% agarose gels, purified and sequenced at the Yale Keck DNA Sequencing facility. New sequences were combined with existing data [47] for a total data set of 490 sequences (S1 Table). For microsatellite analysis, we genotyped flies at 16 microsatellite loci using methods described in [47]. New genotype calls were combined with existing data [47] for a total data set of 558 genotypes at 16 microsatellite loci (S1 Table).

### Identifying *Gff* mtDNA and nuclear genetic backgrounds

To evaluate the potential association between *Spiroplasma* infection prevalence and the *Gff* genetic background, we assigned each individual to a single mtDNA haplogroup based on the phylogenetic relationships among haplotypes, and to a single nuclear genetic background based on clustering analysis of microsatellite genotypes. For mtDNA haplogroup assignment, first evolutionary relationships between the mtDNA haplotypes were assessed by constructing a parsimony-based network using TCS 1.21 [55] as implemented in PopART ([56]; Population Analysis with Reticulate Trees: http://otago.ac.nz). These haplotypes were then grouped by phylogenetic relationship following [48] into three haplogroups, each imperfectly associated with the NWGU, NEGU, and WGU.

For nuclear genetic background assignment, we used a Bayesian clustering analysis in the program STRUCTURE v2.3.4 [57] to group individuals based on their microsatellite genotypes. STRUCTURE assigns individuals into a given number of clusters (K) to maximize Hardy-Weinberg and linkage equilibrium. The program calculates the posterior probability for a range of K and provides a membership coefficient (q-value) of each individual to each cluster. In this analysis, we assessed membership to just three clusters corresponding to the NWGU, NEGU, and WGU. The ADMX does not represent its own cluster, but instead represents a mixed assignment to the NWGU and NEGU. We performed 10 independent runs for K=3 with a burn-in of 50.000 followed by 250.000 MCMC steps and summarized results across the 10 independent MCMC runs using the software CLUMPAK [58]. Individual flies were assigned to nuclear genetic units based on q-values. Individuals with q-values > 0.8 were assigned to one of the three distinct clusters (NWGU, NEGU or WGU), and individuals with mixed assignment (q-values ranging from 0.2 to 0.8) to the NWGU and NEGU were assigned as “admixed” (ADMX).

### Associations of *Spiroplasma* infection with watershed, season of collection, and host genetic traits

Predictive variables considered included watershed of origin, season of collection, *Gff* host genetic background (both nuclear and mitochondrial), *Gff* host sex, and trypanosome co-infection. A challenge in this analysis was that correlation between the geographic origin (specifically watershed) and environmental conditions, as well as *Gff* genetic background and trypanosome infection status is well established [47-49,59]. We confirmed these correlations among predictive variables and patterns of *Spiroplasma* infection, we performed a multiple correspondence analysis (MCA).

We took two approaches to control for correlation among predictive variables with watershed of origin. First, we fit generalized logistic mixed models (GLMM) with the predictive variables of interest (season, *Gff* host genetic background, sex, and co-infection) as fixed effects with and without watershed of origin as the random effect, and tested the improvement of the models with an analysis of variance (ANOVA) Chi square test. We followed these tests with more complex combinations of predictive variables using watershed as the random effect. GLMM was performed with the ‘glmer’ function in the R [60] package *lme4* [61] and fitted using maximum likelihood.

Second, we fit generalized linear models (GLM) one watershed at a time for each predictive variable (season, *Gff* host nuclear and mtDNA genetic background, sex, and co-infection). Direction and significance of the effect of each predictive variable was assessed with Tukey’s contrasts with p-values (*P)* obtained from the z distribution and corrected for multiple comparisons using the unconstrained (“free”) adjustment. GLM was performed in R with the *multcomp* package [62].

### Effect of *Spiroplasma* and trypanosome infection success under laboratory conditions

Effect of *Spiroplasma* presence on trypanosome infection outcome was also tested in the laboratory by infecting the *Gff* colony (IAEA, Vienna, Austria), with bloodstream form *Trypanosoma brucei brucei* (RUMP503) parasites. This *Gff* line has been shown to exhibit a heterogeneous *Spiroplasma* infection prevalence [31]. Following our established protocols [63,64], teneral (newly eclosed) flies were infected by supplementing their first blood meal with 5×10^6^ parasites/ml. All flies that had successfully fed on the infectious blood meal were subsequently maintained on normal blood, which they received every other day. Fourteen days post infection (dpi), the presence of trypanosome infections in the midgut was assessed microscopically plus using a PCR assay. PCR was performed using primers trypalphatubF and trypalphatubR, which target the alpha chain of *T. brucei tubulin* (S2 Table). The reaction set up and the cycler profile were the same as used for *Wolbachia ARM*-PCR described above. *Spiroplasma* presence was assessed in the corresponding reproductive tissue of each fly via PCR assay using the *Spiroplasma* infection assay described above. We used generalized linear models (GLM) to test for the significance of difference in trypanosome infection between *Spiroplasma*-infected and uninfected flies.

## Results

### *Spiroplasma* infection type from field collections

To assess the *Spiroplasma* strain infecting *Gff* sampling sites, we employed 16S *r*DNA and *rpo*B sequencing analysis. In samples analyzed from the Okole River and Lake Kyoga watersheds, we found a single strain infection (S1 Fig), which belongs to the *Citri-Chrysopicola-Mirum* clade and which was also previously identified from a *Gff* colony [31].

We tested for *Spiroplasma* infection in 1415 *Gff* individuals (894 females and 522 males; S1 Table) collected from 26 sampling sites spanning five watersheds (Fig 1). In the northwest region, flies from the Albert Nile watershed (n = 487) had a mean infection rate of 34% (Fig 1; S1 Table). In the northcentral region, flies from the Achwa River watershed (n=234) had a mean infection rate of 20%, and the Okole River watershed (n=389) had a mean infection rate of 5% (Fig 1; S1 Table). In the northeast region, flies from the Lake Kyoga watershed had relatively low *Spiroplasma* infection rate, with one of the two sampling sites (AMI, n = 73) having an infection rate of 11%, and the other site (OCU, n = 90) lacking *Spiroplasma* infection altogether (Fig 1; S1 Table). In the western region, flies from the Kafu River watershed (n = 142) had an infection rate of 3% (Fig 1, S1 Table). These results indicate higher *Spiroplasma* infection in the northwest than in the northeast or west of Uganda, a conclusion that was further supported by tests for association of *Spiroplasma* with watershed of origin using generalized linear modeling (see discussion of MDS, GLMM, and GLM results below).

**Fig 1.**
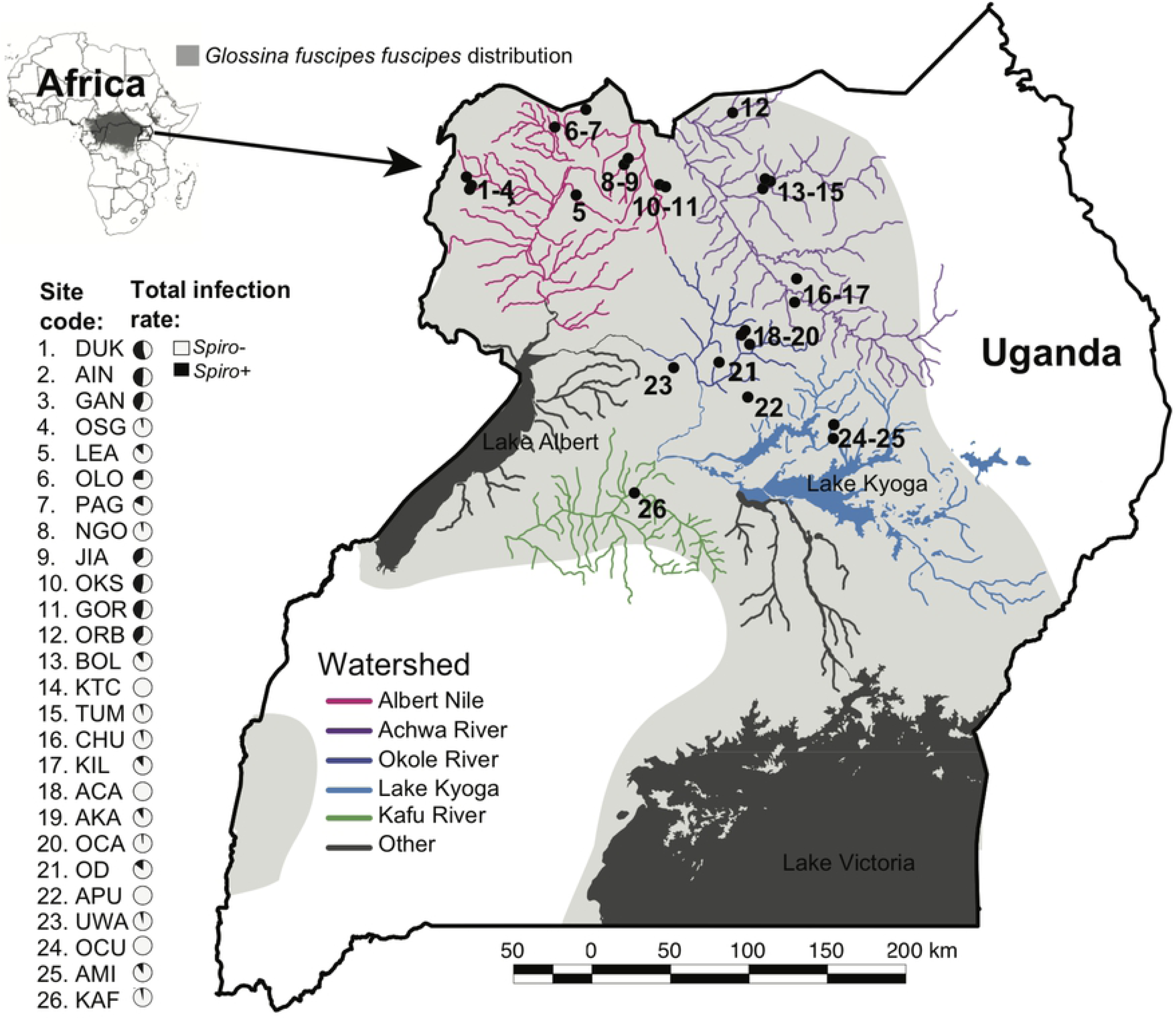
Sampling sites and Spiroplasma infection prevalence in northern Uganda. The map shows the 26 sampling sites and the five watersheds sampled for this study. The location of each sampling site was placed on a map of Uganda using QGIS v2.12.1 (August 2017; http://qgis.osgeo.org) with free and publicly available data from DIVA-GIS (August 2017; http://www.diva-gis.org). Light grey shading shows the distribution of Gff in Uganda. The Spiroplasma infection prevalence is shown in pie charts next to each sampling site (gray = Spiroplasma-positive, black = Spiroplasma-negative).

### Variation in *Spiroplasma* density

To assess whether potential differences in *Spiroplasma* infection density could influence our ability to detect the microorganism in the DNA source, we performed qPCR on individuals from the NEGU (AMI) sampled across multiple seasons. We detected varying densities, with the highest *Spiroplasma* levels observed in the intermediate season (S2 Fig).

### Assignment of *Gff* nuclear and mitochondrial genetic background

We assigned *Gff* host nuclear (biparentally inherited) genetic background using STRUCTURE cluster assignments that were based on the microsatellite genotypes. STRUCTURE assignment indicated 195 flies had high (> 0.8) membership probability to the NWGU, 94 to the NEGU, and 109 to the WGU (S1 Table). We found 160 flies with mixed assignment, which we considered members of the ADMX genetic background (S1 Table). Flies assigned to the NWGU had a mean *Spiroplasma* infection rate of 25%, the ADMX genetic background had a mean infection rate of 15%, the NEGU (n=94) had a mean infection rate of 2%, and the WGU had a mean infection rate of 4% (Fig 2; S1 Table). Although these results suggest higher *Spiroplasma* infection in the NWGU and ADMX nuclear genetic backgrounds, this pattern was found to driven by correlation between nuclear genetic background and watershed of origin (see discussion of MDS, GLMM, and GLM results below). We assigned *Gff* host mitochondrial (maternally inherited) genetic background by generating a TCS network of the mtDNA sequences (n=490). We identified 29 unique mitochondrial haplotypes that grouped into the three previously described major haplogroups: mtA, mtB and mtC (Fig 2, S4 Fig) [47]. 266 flies were assigned to haplogroup A, 209 to haplogroup B, and 15 to haplogroup C (S4 Fig). Flies assigned to mtA had a mean *Spirplasma* infection rate of 19%, mtB had a mean infection rate of 10%, and mtC had a mean infection rate of 13% (Fig 2; S1 Table). Although these results suggest higher *Spiroplasma* infection in the mtA mitochondrial genetic background, this pattern was found to be driven by correlation between mitochondrial genetic background and watershed of origin (see discussion of MDS, GLMM, and GLM results below).

**Fig 2.**
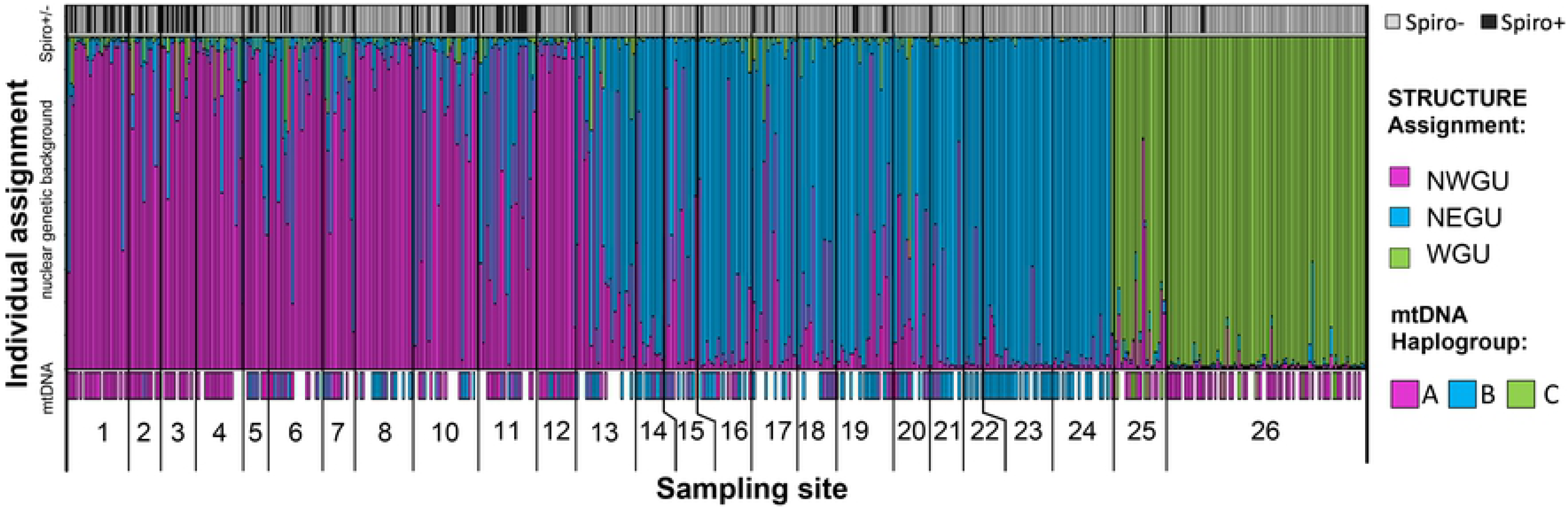
Plot of Gff host Spiroplasma infection, STRUCTURE assignment to nuclear genetic background, and mtDNA haplogroup. Each bar represents a single individual. The top panel shows presence/absence of Spiroplasma of each individual (gray = negative, black = positive). The middle panel shows the STRUCTURE assignment to nuclear genetic background of each individual (pink = northwest genetic unit, blue = northeast genetic unit, green = west genetic unit). The bottom panel shows the mtDNA haplogroup assignment to each individual (pink = mtA, blue = mtB, green = mtC, blank = missing data). Numbers below the bar plot correspond to the 26 sampling sites shown on the map in Fig 1.

### Associations of *Spiroplasma* infection with watershed of origin, season, and host traits under field collections

The MCA confirmed that there were strong correlations among predictive variables, especially with watershed of origin (S3 Fig). GLMM indicated that watershed of origin and season of collection (wet, intermediate and dry) were the two most important factors influencing *Spiroplasma* prevalence (Table 1, S3 and S4 Tables). GLMM of all predictive variables (considered one at a time) were significantly improved by adding watershed as a random effect (ANOVA *P* ranging from 2.20e-16 to 0.0022, S3 Table). Models exploring multiple predictive variables at a time indicated that season of collection was the only variable that positively influenced the fit of the model (ANOVA *P =* 3.57e-6, Table 1). Adding random slope or any of the other predictive variables (nuclear or mtDNA genetic background, sex, or trypanosome co-infection) did not significantly improve the model (ANOVA *P* ranging from 0.2870 to 1.0, S4 Table).

**Table 1.**
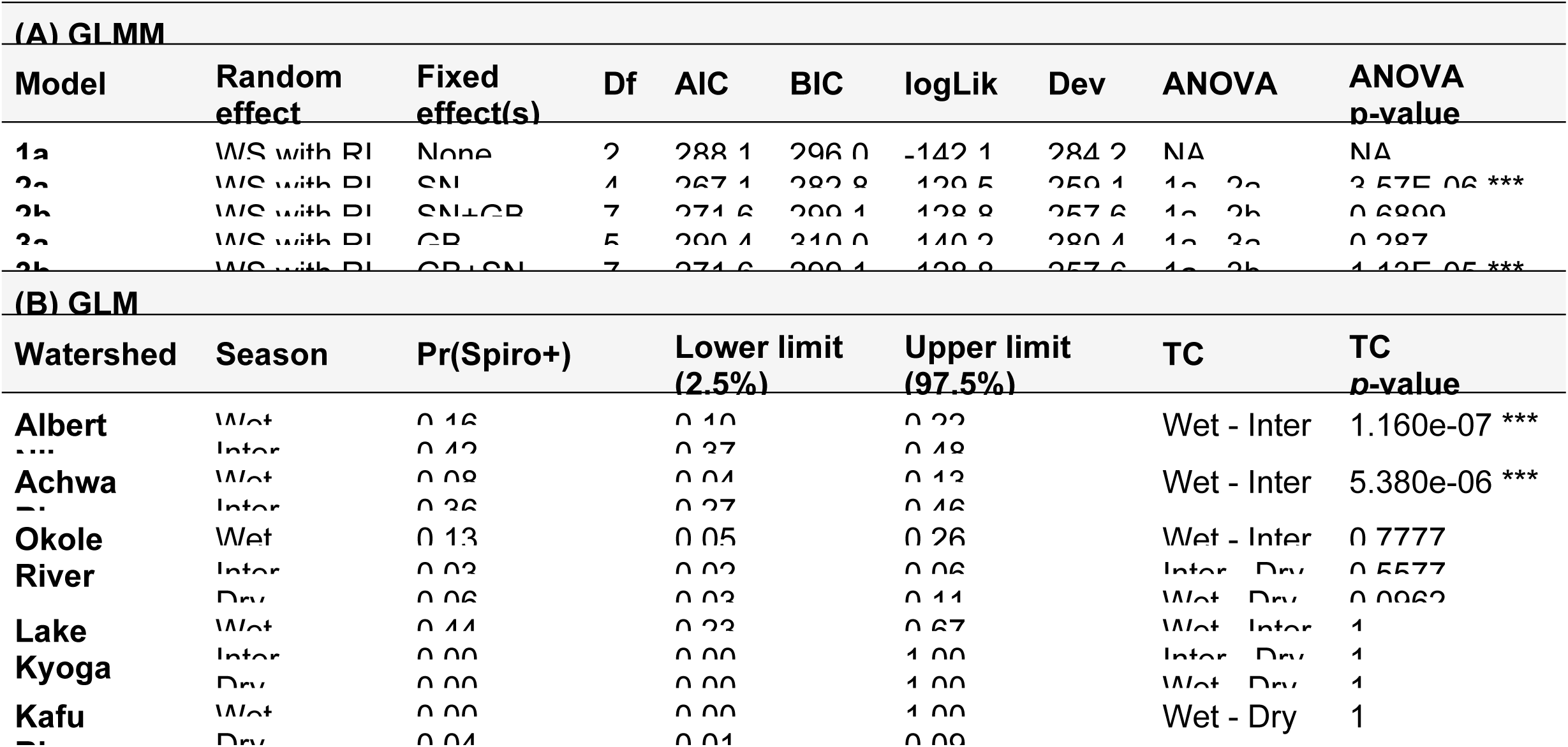
Influence of different predictors on the probability of Spiroplasma infection [Pr(Spiro+)] in generalized linear models. (A) Results from generalized linear mixed models (GLMM), where the random effect was watershed (WS) with random intercept (RI), and the fixed effects were season (SN) and/or nuclear genetic background (GB). The table includes the model name, the fixed effect(s), the degrees of freedom (Df), the Akaike information criterion (AIC) with the best model in bold, the Bayesian information criterion (BIC) with the best model in bold, the log likelihood (logLik), the deviance (Dev) of the model, the analysis of variance (ANOVA) used to test for the model’s improvement, the ANOVA Chi square value (ChiSq), the Chi Square degrees of freedom (ChiDf), and the ANOVA p-value. Adding SN was the only fixed effect that significantly improved the model. In contrast, GB did not significantly improve the GLMM. Additionally, changing the order introducing fixed effects or adding random slope also did not improve the GLMM (S3 Table). (B) Generalized linear model (GLM) by watershed, with season as the predictor. The table includes watershed, season of collection [wet, intermediate (Inter), and dry], GLM estimate of Pr(Spiro+), the lower 2.5% limit and the upper 97.5% limit of the estimate, the Tukey’s contrast (TC) used to test for significant differences in Pr(Spiro+) between seasons, and the TC p-value. The only significant differences in Pr(Spiro+) between seasons was between the wet and the intermediate seasons in the Albert Nile and Achwa River watersheds. Additionally, none of the other predictors (GB, mtDNA haplogroup, trypanosome infection status, or sex) had significant effect on Pr(Sprio) (S5 Table). For all p-values, the level of significance was marked *** if < 0.0001, ** if < 0.001, * if < 0.05, and not marked if > 0.05.

GLM by watershed indicated that flies from the Albert Nile had by far the highest probability of *Spiroplasma* infection [Pr(Spiro+) = 0.32], followed by flies from the neighboring Achwa River [Pr(Spiro+) = 0.22, Fig 3]. Tukey’s contrasts indicated that the Albert Nile had only somewhat higher Pr(Spiro+) than the Achwa River (Tukey’s contrast *P =* 0.0015), but that these two watersheds had significantly higher Pr(Spiro+) than any of the other watersheds (Tukey’s contrasts *P* ranging from 2.00e-16 to 0.0002, S5 Table).

**Fig 3.**
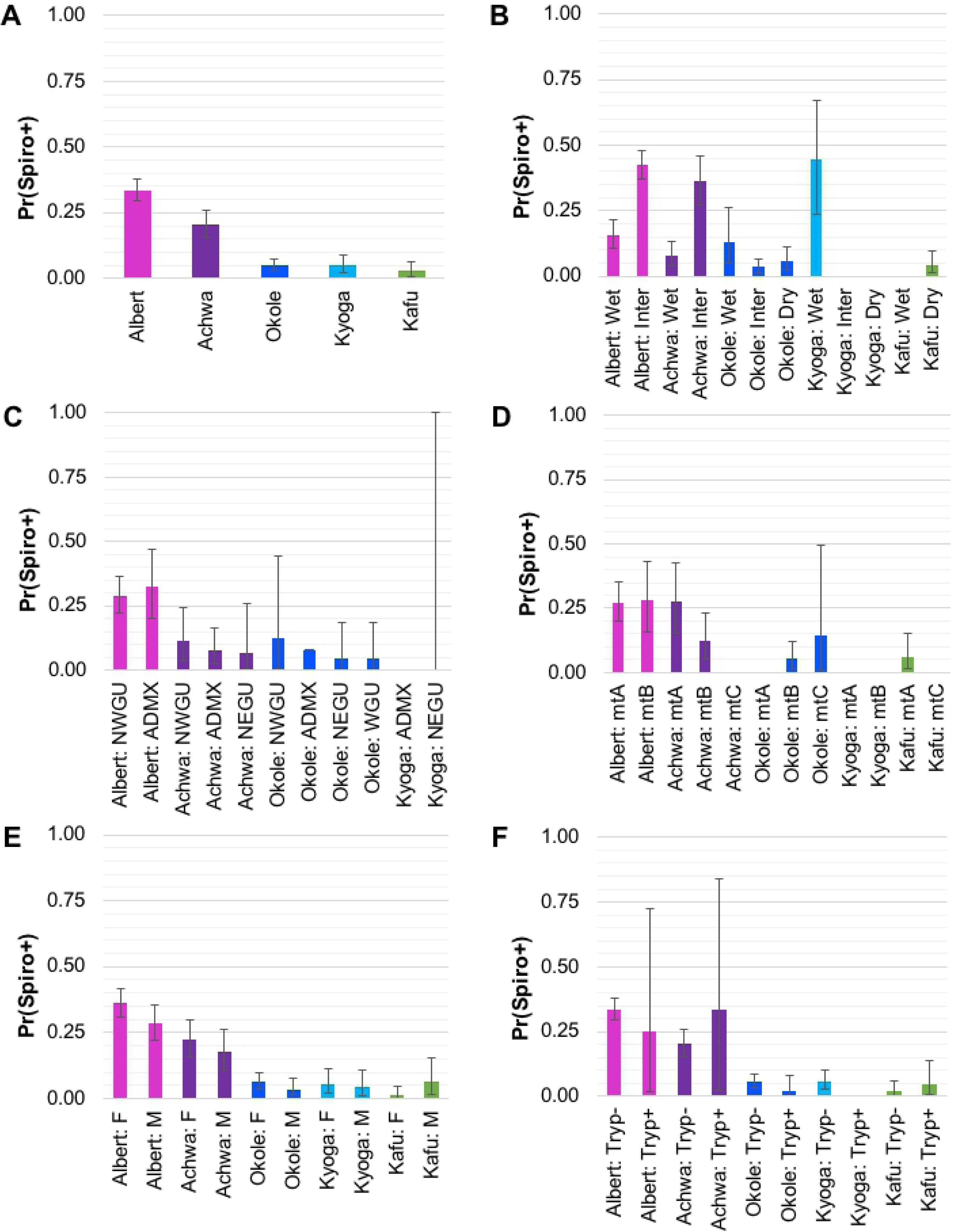
Probability of Spiroplasma infection by watershed. Results of the generalized linear models (GLM) by watershed of origin showing estimated probability of Spiroplasma infection [Pr(Spiro+)] with lower 2.5% and upper 97.5% bounds for (A) watershed of origin only, (B) season of collection by watershed, (C) Gff host nuclear genetic background (NWGU = northwest genetic unit, ADMX = admixed, NEGU = northeast genetic unit, WGU = west genetic unit), (D) Gff host mitochondrial genetic background (mtA = haplogroup A, mtB = haplogroup B, mtC = haplogroup C), (E) Gff host sex (F = female, M = male), and (F) trypanosome coinfection (Tryp+ = coinfection, Tryp-= no coinfection). Each bar is labeled with the watershed and predictive variable, and color coded by watershed (pink = Albert Nile, purple = Achwa River, dark blue = Okole River, light blue = Lake Kyoga, green = Kafu River). See S5 Table for details and Tukey’s contrasts that assess significance of the associations.

In addition to watershed of origin, season of collection was strongly associated with *Spiroplasma* infection. The intermediate season had significantly higher probability of *Spiroplasma* infection than the wet season, which was especially apparent in the Albert Nile [Pr(Spiro+) = 0.42 vs 0.16)] and Achwa River watersheds [Pr(Spiro+) = 0.36 vs 0.08] (Tukey’s contrasts *P* = 1.16E-07 and 5.38E-06, respectively; Table 1). There were no other significant differences in Pr(Spiro+) among seasons in any of the other watersheds (Table 1). Additionally, none of the other predictive variables (nuclear or mtDNA genetic background, sex, or trypanosome co-infection) had significant effects on Pr(Spiro+) when analyzed by watershed (Tukey’s contrasts *P* ranging from 0.0962 to 1.0, S5 Table).

### Co-infections of *Spiroplasma* and *Wolbachia* symbionts

Previous analysis of *Gff* from southern and central regions of Uganda also indicated presence of distinct genetic backgrounds associated with these individuals [47,48] as well as the presence of heterogeneous and low-density infections with another endosymbiont, *Wolbachia* [27, 31]. To investigate for similar patterns in northern Uganda, we analyzed a subset of *Gff* individuals from the Albert Nile (NWGU) and Achwa/Okole River (ADMX) watersheds for the presence of *Wolbachia* infections. We did not detect presence of *Wolbachia* in either *Spiroplasma*-infected (n = 15) or *Spiroplasma*-uninfected individuals (n = 40; S6 Table), which made it impossible to apply any statistical tests to the results.

### Effects of *Spiroplasma* on probability of trypanosome coinfection under laboratory conditions

Patterns of *Spiroplasma* infection from the field collections, although not significant in the GLMM or GLM, indicated a possible correlation between *Spiroplasma* and trypanosome co-infections. Of the 243 *Spiroplasma* infected flies identified in the study, only 2% (n=5) had trypansome co-infection. To test this correlation under more controlled conditions where we could ensure statistical power, we used a laboratory line of *Gff* with heterogeneous infection of *Spiroplasma* [30] to complete a trypanosome infection challenge experiment. Microscopic examination and PCR analysis of challenged individuals (n=94) revealed a negative correlation between the presence of *Spiroplasma* and trypanosome co-infection, with 20% of individuals co-infected with both microbes (*S*^+^*T*^+^; n = 19), 21% infected with only trypanosomes (*S*^-^*T*^+^; n = 20), 43% infected with only *Spiroplasma* (*S*^+^*T*^-^; n = 40), and 16% without infection with either microbe (*S*^-^*T*^-^; n = 15; S6 Table). The generalized mixed models (GLM) indicated a significant negative correlation between the presence of *Spiroplasma* and the probability of trypanosome coinfection (*Tukey’s P* = 0.0192).

## Discussion

In this study, we performed a spatial and temporal analysis of the prevalence of the endosymbiont *Spiroplasma* in genetically distinct *Gff* populations across northern Uganda and assessed the factors that could impact the dynamics of infection with this symbiotic bacterium. The most influential factors that shaped the patchy distribution of *Spiroplasma* in *Gff* were the fly’s watershed of origin and the season of collection. The importance of watershed and season suggests that the prevalence of the symbiont is significantly affected by the geographic dispersal of the flies as well as by the changing environmental conditions. The low rate of *Spiroplasma* and trypanosome co-infections in the field collections, and the negative association of *Spiroplasma* and with trypanosome infection in a challenge experiment carried out in a laboratory *Gff* line suggests that *Spiroplasma* infections may negatively influence the success of parasite infection outcome.

### Geographic origin and seasonality

We found that flies residing in geographically separated watersheds have significantly different *Spiroplasma* infection prevalences (Fig 1, S3 Fig). *Gff* is a riparian species that inhabits low bushes or forests at the margins of rivers, lakes or temporarily flooded scrub land. These water connections may influence dispersal patterns by limiting dispersal between the different watersheds. Limited dispersal would minimize contact among flies from different watersheds. This could suggest that horizontal transfer occurs between flies within the same watersheds, or that there is environmental acquisition of *Spiroplasma* in *Gff*. However, it remains to be elucidated where flies encounter *Spiroplasma* from the environment or during feeding. Interestingly, although tsetse are strict blood feeders on vertebrate hosts, a recent study has indicated that they are capable of feeding on water with or without sugar when deprived of a blood meal [65]. This feeding behavior would allow *Gff* to acquire *Spiroplasma* from the environment, and could account for the strong association of *Spiroplasma* with watershed of origin, and lack of correlation with other host genetic background.

In addition to geographic origin, the season of collection was an important factor shaping the *Gff*-*Spiroplasma* association in Uganda. We found that the intermediate season (June - July) was correlated with higher *Spiroplasma* infection prevalence than either the wet (April - May and August - October) or dry (December - March) seasons (Table 1, Fig 3). This might be because the environmental conditions during the wet and dry seasons restrict the availability of animal hosts for tsetse blood feeding. This could indicate that the intermediate season represents optimal foraging conditions for *Gff*. In fact, *Spiroplasma* survival in *Drosophila* is dependent on the availability of hemolymph lipids [43,44]. Hence, during nutritionally optimal times, maintenance of the symbiont may be less costly for the host than during a period of compromised fitness, and thus higher prevalence and density is observed during these periods [43,44]. Other symbionts capable of triggering host phenotypes, such as male-killing, are also affected by host fitness. The persistence of *Wolbachia*, for example, is negatively affected by host fitness via temperature as well as by other stress factors (e.g. [66-68]). Thus, the more stressful conditions of the dry season may reduce the overall fitness of the fly and consequently the *Spiroplasma* infection densities. This scenario is supported by our finding of different *Spiroplasma* densities in flies analyzed across the three sampling seasons (Fig 2, S2 Fig). Varying *Spiroplasma* densities can also influence the transmission efficiency of the symbiont from mother to tsetse’s intrauterine progeny. Furthermore, if *Spiroplasma* is horizontally transferred via the environment, the climatic conditions can also impact the abundance of *Spiroplasma* for acquisition by *Gff*. Higher *Spiroplasma* density in the environment during the intermediate season could facilitate their acquisition by the tsetse host. In support of this theory, free-living bacterial communities that reside in aquatic systems are strongly affected by environmental factors, such as pH [69,70] and seasonality [71]. A more recent study has highlighted the significant effect of seasonality-related changes in soil bacterial communities [72]. Hence, the changing environmental conditions that occur between the dry, wet, and intermediate season might impact *Spiroplasma* density and consequently the infection density.

### Host genetic nuclear and mitochondrial background

Genetically variable *Gff* populations residing within the Gambiense and Rhodesiense HAT belts could influence the inheritable microbiota composition, which in turn could have important implications in transmission dynamics and vector control outcomes [73- 75]. In that context, we evaluated the potential correlation between host nuclear genetic differences and *Spiroplasma* infection prevalence across the 26 *Gff* sampling sites. However, we found that after accounting for watershed of origin in the GLMM model, *Spiroplasma* infection was not significantly influenced by *Gff* host nuclear or mitochondrial genetic background (Figs 2 and 3, S5 Table). Lack of association of with host genetic background provides further support for the idea that *Spiroplasma* may be acquired by horizontal transfer between flies within the same watersheds, or through contact with other sources of the bacteria in the environment.

### *Spiroplasma* and *Wolbachia*

In *Drosophila, Spiroplasma* can either coexist with the endosymbiotic *Wolbachia* in a synergistic manner [76], or negatively affect *Wolbachia* presence. Goto and coworkers showed that the presence of *Spiroplasma* suppresses *Wolbachia* infection density in *D. melanogaster* [77]. Interestingly, while the tsetse species in the *Morsitans* subgroup, such as *G. morsitans*, harbor *Wolbachia* infections, they lack *Spiroplasma* associations [31]. In contrast, the tsetse species in the *Palpalis* subgroup, such as *Gff* and *G. palpalis*, harbor *Spiroplasma* infections, but lack *Wolbachia* associations [31]. *Spiroplasma* might be negatively affected by the presence of *Wolbachia* infection in the species within the *Morsitans* subgroup [31]. As such, the high titer of *Wolbachia* noted in *G. morsitans* reproductive organs could suppress the establishment of *Spiroplasma* infections. Our prior and current studies with distinct *Gff* populations in Uganda also indicate a potential negative influence for the two endosymbiont infections [28]. Our previous finding reported *Wolbachia* infections across 18 *Gff* sampling sites from the Northcentral, West and South of Uganda [28]. In contrast, we did not detect *Wolbachia* infections in the current samples analyzed from the Albert Nile or Achwa River watersheds, which were the geographic areas with highest *Spiroplasma* infection prevalence (S1 Table). Given that *Spiroplasma* and *Wolbachia* infections appear to be associated with geographically and genetically distinct *Gff* populations, the idea of reciprocal exclusion of both entities remains a possibility. However, we had also noted that the *Wolbachia* density in the reproductive organs of *Gff* is unusually low, and the *Spiroplasma* densities can vary across seasons, thus rendering co-infection detections technically challenging, particularly when using whole body DNA, as was the case with a number of flies analyzed in this study. It also remains to be elucidated if the presence of viral microorganisms plays a role for the infection dynamics of *Spiroplasma* in Ugandan *Gff* populations. The salivary gland hypertrophy virus (SGHV), a common virus of *Glossina*, is inversely correlated with *Wolbachia* infection prevalence in *Gff* in Uganda [28]. Hence, the correlation between *Spiroplasma* and SGHV remains to be tested.

### Role of *Spiroplasma* association with *Gff* physiology

Reproductive influences, such as male killing, have been associated with maternally inherited symbionts, including *Spiroplasma,* where the symbiont drives its own dispersal by selectively killing male embryos in ladybirds, fruit flies and certain butterfly species (reviewed in [41]). As we observed both *Spiroplasma*-infected males and females, this bacterium likely does not confer a male killing trait to *Gff*. Pairwise comparison suggested a slightly higher infection prevalence in females, but such a slight difference is unlikely to be the result of a male killing phenotype.

We also evaluated the potential role of *Spiroplasma* on tsetse’s immune physiology by measuring the correlation between trypanosome and *Spiroplasma* infection status. Of the 1415 samples analyzed, we found only five with trypanosome and *Spiroplasma* co-infections. Although not significant in the GLMM or GLM, this negative results could have been caused by lack of statistical power in the field collected data. To further address the question of a similar protective effect, we performed trypanosome infection experiments using a colonized *Gff* line that displays heterogeneous *Spiroplasma* infection prevalence [31]. Our finding that infections with *Spiroplasma* alone were more frequent than co-infections with *Tbb* (43% vs. 20%, respectively; S4 Table) indicates a negative correlation between the presence of the symbiont and the parasite, and suggests that both entities negatively impact each other’s fitness. In accordance with investigations in *Drosophila* [78,79], *Spiroplasma* infections may confer physiological traits that protect its tsetse host from being colonized by trypanosome infections. In different hosts, *Spiroplasma* induces a protective effect against nematodes, fungi and parasitoid wasps [35,36,78]. It remains to be elucidated, whether a potential protection in *Gff* against parasite infections results from niche competition of both microbes within the host, or by expression of certain *Spiroplasma*-derived molecules, which block trypanosomes. Alternatively, it has been shown that infections with certain *Wolbachia* strains confer an immune enhancement phenotype to their *Drosophila* and mosquito hosts, hence increasing the resistance to other pathogens [80; also reviewed in 81]. Such an immune enhancement, which affect the trypanosome transmission success in *Spiroplasma* infected *Gffs* however, is rather unlikely given the lack of *Wolbachia* in the tested flies.

The heterogeneous *Spiroplasma* infection prevalence in the *Gff* line that we used in the parasite challenge experiment (Fig 4) might result from imperfect maternal transmission to progeny and may similarly be influencing the infection prevalence noted in natural populations. Tsetse are viviparous and the mother supports the development of her progeny in an intrauterine environment. Endosymbiotic *Wigglesworthia* and *Sodalis* are maternally acquired by the progeny in female milk secretions during the lactation process, while *Wolbachia* is transovarially transmitted. *Spiroplasma* infections in the gonads suggest that this endosymbiont is also transovarially transmitted, although transmission through milk secretions remains to be investigated. While tsetse females remain fecund throughout their entire life (and can produce 8-10 progeny), the transmission of *Spiroplasma* from mother to her intrauterine progeny however may be more efficient during the early gonotrophic cycles and decrease over the course of female’s reproductive lifespan. Such a scenario could explain why we observed that only 50 −60% of colony flies are infected with *Spiroplasma*. We will further address this question by testing the efficiency of symbiont transmission from mother to each of her offspring in a follow-up study by developing single lines from each pregnant female. Such an imperfect transmission efficiency can also influence the heterogeneous infection prevalence we noted in field populations.

**Fig 4.**
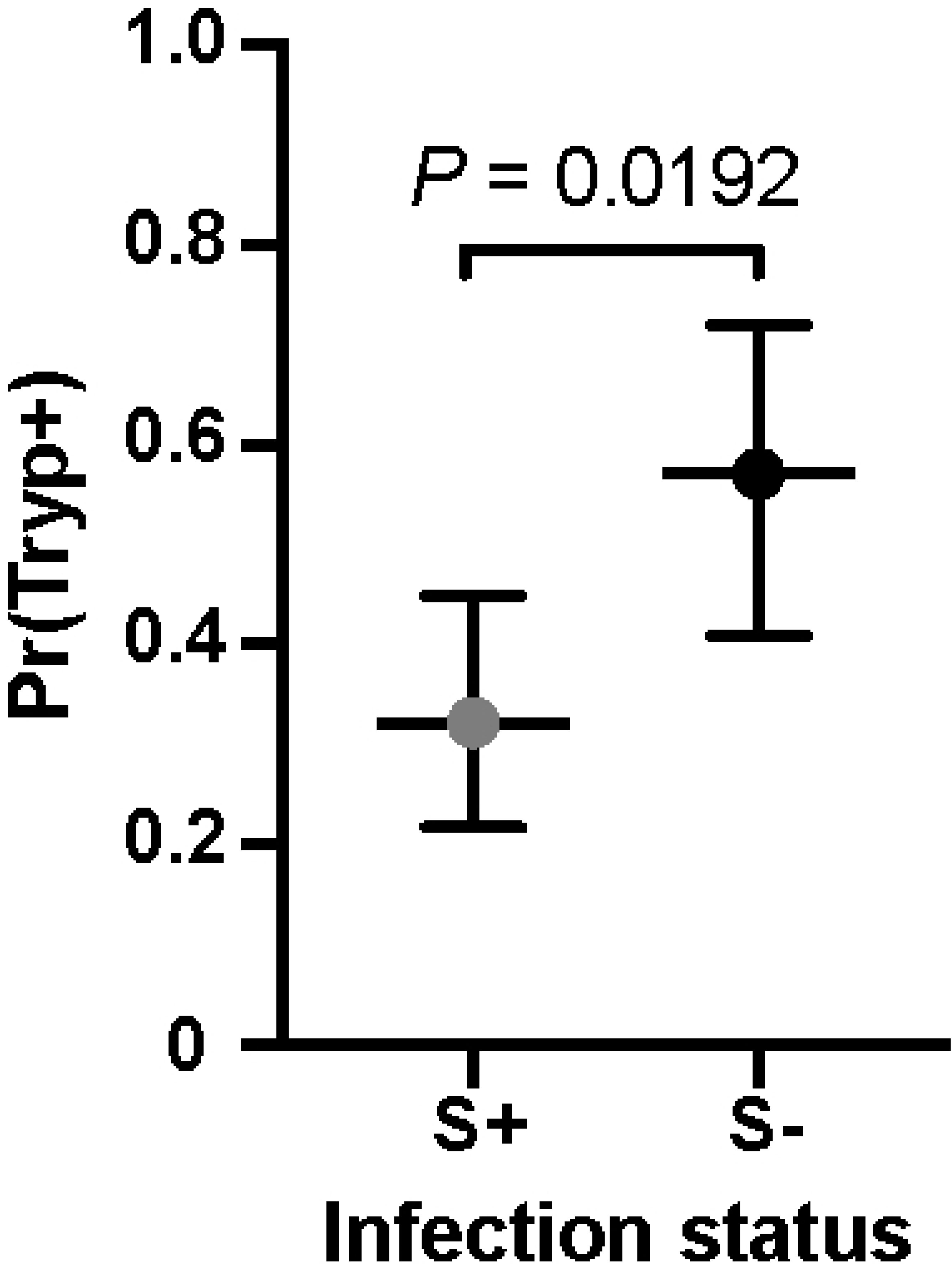
Trypanosome infection experiment using a laboratory Gff line. Scatter dot plot shows the percentage of flies 14 days post infection acquisition found to be infected with either Spiroplasma only (S^+^T^-^, gray) or with with trypanosome only (S^-^T^+^, black). Bars show 95% confidence interval. Infection status was assessed by microscopy (trypanosomes) and PCR (trypanosomes and Spiroplasma). Dotted line shows the median of the data. Abbreviations: S^+^ Spiroplasma-infected, S^-^ Spiroplasma-uninfected, T^+^ Trypanosoma-infected, T^-^ Trypanosoma-uninfected.

In conclusion, the infection prevalence of *Spiroplasma* in *Gff* populations in northern Uganda is significantly correlated with different watersheds of origin and seasonal environmental conditions. These associations indicate that seasonal fluctuations and other transmission modes than strictly vertically are drivers of *Spiroplasma* acquisition in *Gff* in Uganda.

We further demonstrate that colonized *Gff* are less likely to establish trypanosome parasite infections when carrying *Spiroplasma* infections, which is of particular interest in the context of alternative vector control approaches to control trypanosome infections.

## Acknowledgments

We thank the Gulu University field team for sampling *Gff* in Uganda. We also thank Andrew G. Parker and Adly M.M. Abd-Alla from the Insect Pest Control Laboratory, Joint FAO/IAEA Division of Nuclear Techniques in Food and Agriculture, Vienna, Austria for providing *Gff* flies.

## Supporting information

**S1 Fig. Neighbor joining tree based on *Spiroplasma RpoB* sequences.** *Spiroplasma* strains from the Albert Nile watershed (OKS) and the Okole River watershed (ACA) are identical to each other and highly conserved in comparison to the *Spiroplasma* present in two individuals screened from a *Gff* laboratory line (KX159379_sGfus_BL, KX159380_sGfus_SL) and one Uganda field sample (KX159381_sGfus_U350). These sequences were recently published in [31]. The tree was calculated in CLC using Jukes-Cantor as distance measure with 1000 replicates, and rooted against the outgroup *Spiroplasma culicicola*. Scale is 0.3 substitutions per site.

**S2 Fig. *Spiroplasma* density differences in *Gff***. **(A)** The graph shows the relative density of *Spiroplasma* tested via *16S r*DNA in low titer and uninfected *Gff* from the Bole (BOL) population in the Achwa watershed. Flies with high loads of *Spiroplasma* are not included in this graph. Some females and males show variation in their *Spiroplasma* load, *i.e.* they are neither negative not high titer but intermediate. **(B)** Relative density of *Spiroplasma* in *Gff* from AMI population (Lake Kyoga) across wet, dry and intermediate season. *Spiroplasma* levels are higher in intermediate and wet season compared to the dry season.

**S3 Fig. Multiple correspondence analysis**. Dimensions 1 and 2 (accounting for 31.7% and 15.6% of the variation) are plotted. Populations are depicted by triangles. Association with trypanosomes and *Spiroplasma* is shown as open triangles (infected and uninfected). Further associations shown in the plot are the host mtDNA genetic background (mtA, mtB, mtC), the nuclear genetic background (NWGU, ADMX, NEGU, WGU), the season (Dry, Inter, Wet), and the watershed of origin (Albert Nile, Achwa River, Okole River, Lake Kyoga, Kafu River).

**S4 Fig. mtDNA haplogroup network of *Gff COI* gene and correlation of *Spiroplasma* infection to host genotype.** The figure shows the TCS network of the COI gene and the proportion of infected individuals within the mtDNA haplogroups A, B, and C. Haplogroup A is associated with the NWGU, haplogroup B is associated with the NEGU, and haplogroup C is associated with the WGU. Red represents individuals with *Spiroplasma* infection while black represents the uninfected ones. The circle size represents the number of individuals sharing a haplotype; small black nodes represent inferred haplotypes; dashes between haplotypes represent a single mutational step. The haplotype network was generated using the TCS method [55] available in POPART [56].

**S1 Table. Detailed information of *Gff* samples analyzed in this study.** The table on the main tab (‘Individuals’) includes information by individual: Fly ID, sampling site code, sampling site name, district, latitude, longitude, watershed of origin, year of collection, month of collection, season of collection, sex, the tissue used to screen for Spiroplasma (RP=reproductive, WB=whole body), the mitochondrial haplogroup assigned (mtA, mtB, mtC), the STRUCTURE assigned nuclear genetic background (NWGU, ADMX, NEGU, WGU), the PCR-based *Spiroplasma* infection status, the visually-based trypanosome infection status, and the source of the genetic data, with individuals that were genotyped for this study indicated as “this study”. A summary of numbers (totals, females, males, *Spiroplasma* infection) per sampling site and per watershed is included in a separate tab of the same file labeled ‘Sampling Sites’.

**S2 Table. List of primer sets employed in this study.** Sequence of primers for non-quantitative plus quantitative PCR is provided in this list.

**S3 Table. *Spiroplasma* modeling results.** Output from generalized linear mixed models (GLMM) with one predictive variable at a time (with and without watershed as a random effect).

**S4 Table. *Spiroplasma* modeling results.** Output from generalized linear mixed model (GLMM) with multiple predictive variables.

**S5 Table. *Spiroplasma* modeling results.** Output from generalized linear models (GLM) by watershed.

**S6 Table. Details of *Tbb*-challenging experiment.** The table summarizes numbers of analyzed flies upon challenge with *Tbb* and their infection status: *S*^+^*T*^+^, *S*^+^*T*^*-*^, *S*^-^*T*^+^, *S*^-^*T*^-^. All flies were analyzed 14 days post infection. Infection status by individual is given in an extra sheet (by individual). The GLM sheet includes the details if the GLM, which was run to analyze the significance of the difference between groups. Abbreviations: *Tbb Trypanosoma brucei brucei*. *S*^+^ *Spiroplasma*-infected, *S*^-^ *Spiroplasma*-uninfected, *T*^+^ *Tbb*-infected, *T*^-^*Tbb*-uninfected.

